# Genome-wide analysis of self-reported risk-taking behaviour and cross-disorder genetic correlations in the UK Biobank cohort

**DOI:** 10.1101/177014

**Authors:** Rona J. Strawbridge, Joey Ward, Breda Cullen, Elizabeth M. Tunbridge, Sarah Hartz, Laura Bierut, Amy Horton, Mark E. S. Bailey, Nicholas Graham, Amy Ferguson, Donald M. Lyall, Daniel Mackay, Laura M. Pidgeon, Jonathan Cavanagh, Jill P. Pell, Michael O’Donovan, Valentina Escott-Price, Paul J. Harrison, Daniel J. Smith

**Affiliations:** Institute of Health and Wellbeing, University of Glasgow, Glasgow, UK; Department of Medicine Solna, Karolinska Institute, Stockholm, Sweden; Department of Psychiatry, University of Oxford, Oxford, UK; Oxford Health NHS Foundation Trust, Oxford, UK; Department of Psychiatry, Washington University School of Medicine in St. Louis, St. Louis, Missouri, USA; Transmontane Analytics, Tuscon, Arizona, USA; School of Life Sciences, College of Medical, Veterinary and Life Sciences, University of Glasgow, Glasgow, UK; MRC Centre for Neuropsychiatric Genetics and Genomics, Cardiff University, Cardiff, UK

## Abstract

Risk-taking behaviour is a key component of several psychiatric disorders and could influence lifestyle choices such as smoking, alcohol use and diet. As a phenotype, risk-taking behaviour therefore fits within a Research Domain Criteria (RDoC) approach, whereby identifying genetic determinants of this trait has the potential to improve our understanding across different psychiatric disorders. Here we report a genome wide association study in 116 255 UK Biobank participants who responded yes/no to the question “Would you consider yourself a risk-taker?” Risk-takers (compared to controls) were more likely to be men, smokers and have a history of psychiatric disorder. Genetic loci associated with risk-taking behaviour were identified on chromosomes 3 (rs13084531) and 6 (rs9379971). The effects of both lead SNPs were comparable between men and women. The chromosome 3 locus highlights *CADM2*, previously implicated in cognitive and executive functions, but the chromosome 6 locus is challenging to interpret due to the complexity of the HLA region. Risk-taking behaviour shared significant genetic risk with schizophrenia, bipolar disorder, attention deficit hyperactivity disorder and post-traumatic stress disorder, as well as with smoking and total obesity. Despite being based on only a single question, this study furthers our understanding of the biology of risk-taking behaviour, a trait which has a major impact on a range of common physical and mental health disorders.

## Introduction

Risk-taking behaviour is an important aspect of several psychiatric disorders, including attention deficit hyperactivity disorder (ADHD) ^1, 2^ and bipolar disorder (BD) ^3^, as well as problem behaviours such as smoking and drug and alcohol misuse ^4, 5^. The link between risk-taking behaviour and schizophrenia (SCZ) is more complex, with difficulties in conditional reasoning ^6^, problems with delayed gratification and poor impulse control occurring alongside more conservative risk assessment ^7^. Physical health problems such as obesity might also be considered to be related to increased propensity towards risk-taking: obesity includes aspects of aberrant reward processing, response inhibition and decision-making ^8^. The Research Domain Criteria (RDoC) approach suggests that studying dimensional psychopathological traits (rather than discrete diagnostic categories), as well as relevant traits across the whole spectrum (“normal” through to pathological) of the population may be a more useful strategy for identifying biology which cuts across psychiatric diagnoses ^9^. In this respect, risk-taking behaviour is an important phenotype for investigation. It may also be useful for investigating the overlap between psychiatric disorders and conditions such as obesity and smoking.

To date, an association between a locus on chromosome 3 and risk-taking behaviour has been published ^10, 11^, but no genome-wide genetic study with a primary focus on risk-taking behaviour has been conducted. GWAS of related phenotypes, such as impulsivity and behavioural disinhibition, have so far been underpowered for detecting associations at a genome-wide level. Here we conduct a primary GWAS of self-reported risk-taking behaviour in 116 255 participants from the UK Biobank cohort. We use expression quantitative trait loci analysis to highlight plausible candidate genes and we assess the extent to which there is a genetic correlation between risk-taking and several mental and physical health disorders, including ADHD, SCZ, BD, major depressive disorder (MDD), anxiety, post-traumatic stress disorder (PTSD), smoking status (ever smoker), lifetime cannabis use, fluid intelligence, years of education, obesity and alcohol use disorder.

## Materials and methods

### Sample

UK Biobank is a large population cohort which aims to investigate a diverse range of factors influencing risk of diseases which are common in middle and older age. Between 2006 and 2010, more than 502 000 participants (age range from 40 and 69 years) were recruited from 22 centres across the UK ^12^. Comprehensive baseline assessments included social circumstances, cognitive abilities, lifestyle and measures of physical health status. The present study used the first release of genetic data on approximately one third of the UK Biobank cohort. In order to maximise homogeneity, we included only participants of (self-reported) white United Kingdom (UK) ancestry.

Informed consent was obtained by UK Biobank from all participants. This study was carried out under the generic approval from the NHS National Research Ethics Service (approval letter dated 13 May 2016, Ref 16/NW/0274) and under UK Biobank approval for application #6553 “Genome-wide association studies of mental health” (PI Daniel Smith).

### Genotyping, imputation and quality control

The first release of genotypic data from UK Biobank, in June 2015, included 152 729 UK Biobank participants. Samples were genotyped with either the Affymetrix UK Biobank Axiom array (Santa Clara, CA, USA; approximately 67%) or the Affymetrix UK BiLEVE Axiom array (33%), which share at least 95% of content. Autosomal data only were available.

Imputation of the data has previously been described in the UK Biobank interim release documentation ^13^. In brief, SNPs were excluded prior to imputation if they were multiallelic or had minor allele frequency (MAF) <1%. A modified version of SHAPEIT2 was used for phasing and IMPUTE2 (implemented on a C++ platform) was used for the imputation ^14, 15^. A merged reference panel of 87 696 888 biallelic variants on 12 570 haplotypes constituted from the 1000 Genomes Phase 3 and UK10K haplotype panels ^16^ was used as the basis for the imputation. Imputed variants with MAF <0.001% were filtered out of the dataset used for subsequent analysis.

The Wellcome Trust Centre for Human Genetics applied stringent quality control, as described in UK Biobank documentation ^17^, before release of the genotypic data set. UK Biobank genomic analysis exclusions were applied (Biobank Data Dictionary item #22010). Participants were excluded from analyses due to relatedness (#22012: genetic relatedness factor; one member of each set of individuals with KING-estimated kinship coefficient >0.0442 was removed at random), sex mismatch (reported compared to genetic) (#22001: genetic sex), non-Caucasian ancestry (#22006: ethnic grouping; self-reported and based on principal component (PC) analysis of genetic data), and quality control failure (#22050: UK BiLEVE Affymetrix quality control for samples and #22051: UK BiLEVE genotype quality control for samples). SNPs were removed due to deviation from Hardy–Weinberg equilibrium at *P*<1×10^−6^, MAF <0.01, imputation quality score <0.4 and >10% missingness in the sample after excluding genotype calls made with <90% posterior probability.

The second release of genetic data from the UK Biobank (July 2017) included a further 349 935 samples. Genotyping platforms, quality control and pre-imputation procedures were consistent with the first data release. Imputation of genotypes at additional SNP loci for all participants (n=502 664) was carried out using the Haplotype Reference Consortium reference panel, and post-imputation quality control was consistent with that of the first data release.

### Risk-taking phenotype

The baseline assessment (2006-2010) of UK Biobank participants included the question “Would you describe yourself as someone who takes risks?” (data field #2040), to which participants replied yes or no. Individuals who responded ‘yes’ to the risk-taking question are here referred to as ‘risk-takers’ and those who responded ‘no’ are here referred to as not risk-takers or controls. For a subset of participants, the same question (“Would you describe yourself as someone who takes risks?”) was asked at follow-up (2012-2013), enabling an assessment of response consistency.

### Discovery analyses

A total of 116 255 individuals and 8 781 003 variants (first data release) were included in the discovery analysis. 29 703 participants were classed as risk-takers and 86 552 were controls. Association analysis was conducted in PLINK ^18^ using logistic regression, assuming a model of additive allelic effects and models were adjusted for sex, age, genotyping array, and the first 8 genetic PCs (Biobank Data Dictionary items #22009.01 to #22009.08) to control for hidden population stratification. The threshold for GWAS significance was set at p<5×10^−8^. Demographics of the discovery sample set are presented in Table 1. For quality control purposes, a GWAS of the individuals included in the discovery analysis was run with the second release genetic data (HPC-imputed) and using the updated genetic exclusions and covariates used. Using the updated exclusions resulted in a slight increase in the number of individuals included in the analysis: n= 117 755, of whom n= 30 013 were risk-takers and n= 87 742 were non risk-takers. The sex distribution and demographics of this dataset were comparable with those included in the discovery analysis based on the first genetic release (Supplementary Table 1).

**Table 1.**
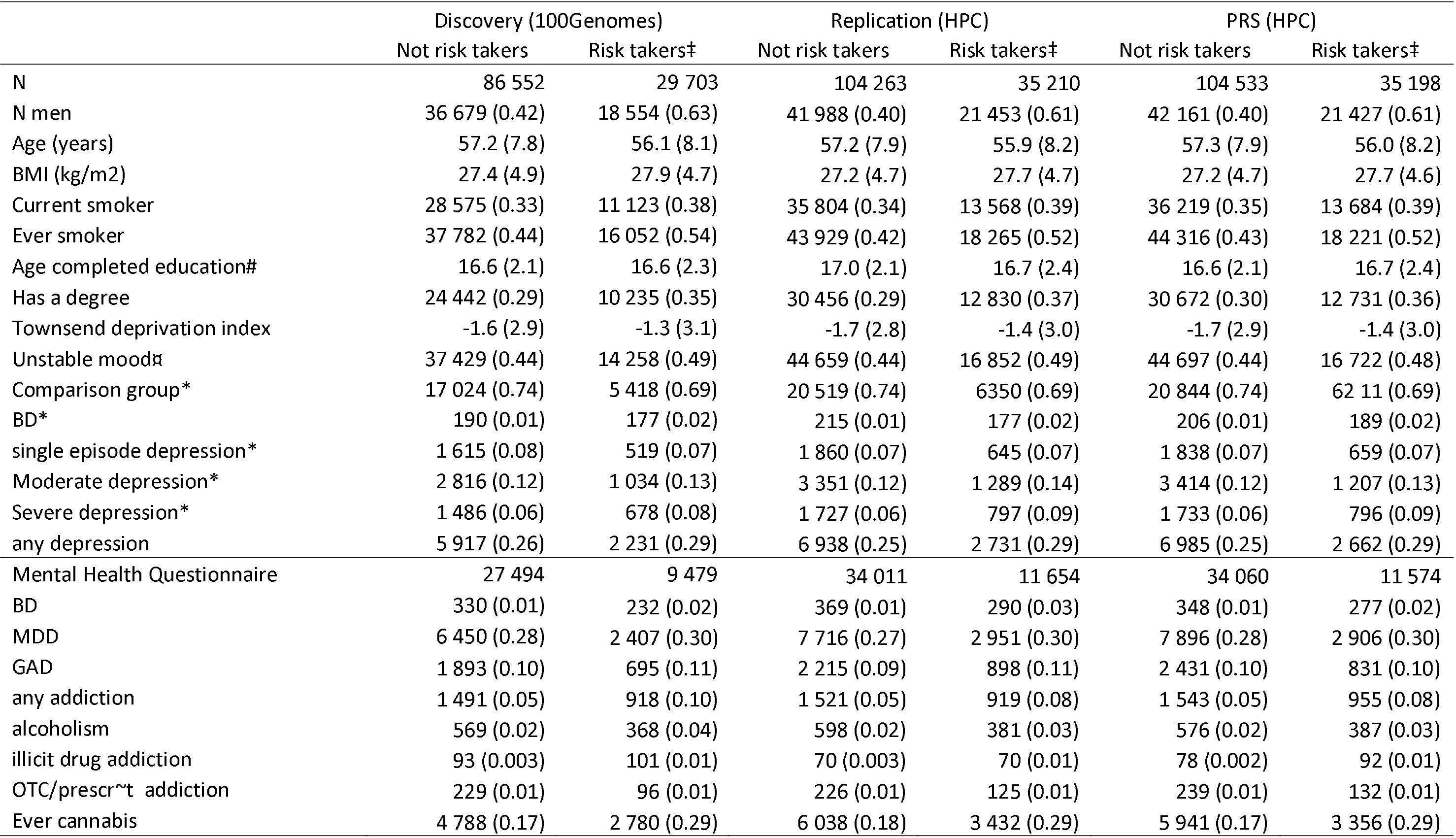
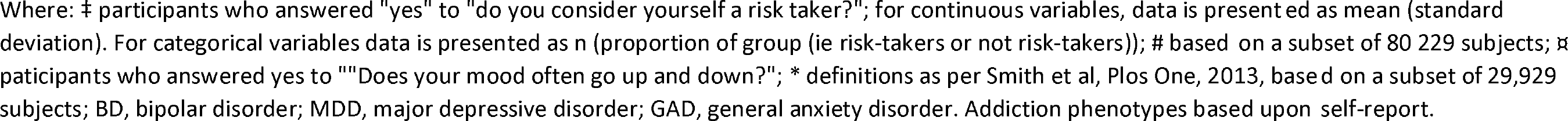
Description of UK Biobank participants included in the discovery risk-taking GWAS, replication and PRS analyses.

|  | Discovery (100Genomes) |  | Replication (HPC) |  | PRS (HPC) |  |
| --- | --- | --- | --- | --- | --- | --- |
|  | Not risk takers | Risk takers‡ | Not risk takers | Risk takers‡ | Not risk takers | Risk takers‡ |
| N | 86 552 | 29 703 | 104 263 | 35 210 | 104 533 | 35 198 |
| N men | 36 679 (0.42) | 18 554 (0.63) | 41 988 (0.40) | 21 453 (0.61) | 42 161 (0.40) | 21 427 (0.61) |
| Age (years) | 57.2 (7.8) | 56.1 (8.1) | 57.2 (7.9) | 55.9 (8.2) | 57.3 (7.9) | 56.0 (8.2) |
| BMI (kg/m<sup>2</sup>) | 27.4 (4.9) | 27.9 (4.7) | 27.2 (4.7) | 27.7 (4.7) | 27.2 (4.7) | 27.7 (4.6) |
| Current smoker | 28 575 (0.33) | 11 123 (0.38) | 35 804 (0.34) | 13 568 (0.39) | 36 219 (0.35) | 13 684 (0.39) |
| Ever smoker | 37 782 (0.44) | 16 052 (0.54) | 43 929 (0.42) | 18 265 (0.52) | 44 316 (0.43) | 18 221 (0.52) |
| Age completed education# | 16.6 (2.1) | 16.6 (2.3) | 17.0 (2.1) | 16.7 (2.4) | 16.6 (2.1) | 16.7 (2.4) |
| Has a degree | 24 442 (0.29) | 10 235 (0.35) | 30 456 (0.29) | 12 830 (0.37) | 30 672 (0.30) | 12 731 (0.36) |
| Townsend deprivation index | -1.6 (2.9) | -1.3 (3.1) | -1.7 (2.8) | -1.4 (3.0) | -1.7 (2.9) | -1.4 (3.0) |
| Unstable moodα | 37 429 (0.44) | 14 258 (0.49) | 44 659 (0.44) | 16 852 (0.49) | 44 697 (0.44) | 16 722 (0.48) |
| Comparison group* | 17 024 (0.74) | 5 418 (0.69) | 20 519 (0.74) | 6350 (0.69) | 20 844 (0.74) | 62 11 (0.69) |
| BD* | 190 (0.01) | 177 (0.02) | 215 (0.01) | 177 (0.02) | 206 (0.01) | 189 (0.02) |
| single episode depression* | 1 615 (0.08) | 519 (0.07) | 1 860 (0.07) | 645 (0.07) | 1 838 (0.07) | 659 (0.07) |
| Moderate depression* | 2 816 (0.12) | 1 034 (0.13) | 3 351 (0.12) | 1 289 (0.14) | 3 414 (0.12) | 1 207 (0.13) |
| Severe depression* | 1 486 (0.06) | 678 (0.08) | 1 727 (0.06) | 797 (0.09) | 1 733 (0.06) | 796 (0.09) |
| any depression | 5 917 (0.26) | 2 231 (0.29) | 6 938 (0.25) | 2 731 (0.29) | 6 985 (0.25) | 2 662 (0.29) |
| Mental Health Questionnaire | 27 494 | 9 479 | 34 011 | 11 654 | 34 060 | 11 574 |
| BD | 330 (0.01) | 232 (0.02) | 369 (0.01) | 290 (0.03) | 348 (0.01) | 277 (0.02) |
| MDD | 6 450 (0.28) | 2 407 (0.30) | 7 716 (0.27) | 2 951 (0.30) | 7 896 (0.28) | 2 906 (0.30) |
| GAD | 1 893 (0.10) | 695 (0.11) | 2 215 (0.09) | 898 (0.11) | 2 431 (0.10) | 831 (0.10) |
| any addiction | 1 491 (0.05) | 918 (0.10) | 1 521 (0.05) | 919 (0.08) | 1 543 (0.05) | 955 (0.08) |
| alcoholism | 569 (0.02) | 368 (0.04) | 598 (0.02) | 381 (0.03) | 576 (0.02) | 387 (0.03) |
| illicit drug addiction | 93 (0.003) | 101 (0.01) | 70 (0.003) | 70 (0.01) | 78 (0.002) | 92 (0.01) |
| OTC/prescr~t addiction | 229 (0.01) | 96 (0.01) | 226 (0.01) | 125 (0.01) | 239 (0.01) | 132 (0.01) |
| Ever cannabis | 4 788 (0.17) | 2 780 (0.29) | 6 038 (0.18) | 3 432 (0.29) | 5 941 (0.17) | 3 356 (0.29) |

Where: ‡ participants who answered "yes" to "do you consider yourself a risk taker?"; for continuous variables, data is presented as mean (standard deviation). For categorical variables data is presented as n (proportion of group (ie risk-takers or not risk-takers)); # based on a subset of 80 229 subjects; ¶ participants who answered yes to ""Does your mood often go up and down?"; \* definitions as per Smith et al, Plos One, 2013, based on a subset of 29,929 subjects; BD, bipolar disorder; MDD, major depressive disorder; GAD, general anxiety disorder. Addiction phenotypes based upon self-report.

### Replication analysis

Approximately half of the participants only present in the second data release were included in the replication analysis, thus after quality control and recommended exclusions, 139 474 white British participants were included. Demographics of the replication sample set are presented in Table 1 The lead SNPs in the *CADM2* and Chr6 loci were selected for replication. Consistent with the discovery analysis, replication analysis was conducted in PLINK ^18^ using logistic regression, assuming a model of additive allelic effects and models were adjusted for sex, age, genotyping array, and the first 8 genetic PCs (PCA1-8) to control for hidden population stratification. As two SNP were investigated, p<0.025 was considered significant. Results were meta-analysed using METAL ^19^.

### Polygenic risk scores

In order to assess the variance explained by the genetic loci identified here, polygenic risk scores (PRS) were calculated in the remaining 50% of the second genetic data release. Demographics of the PRS sample set are presented in Table 1. After quality control and recommended exclusions, 139 731 white British participants were included in this analysis.

PRS were calculated using p-value thresholds of p<1×10^−5^, p<0.001 and p<0.05. A score of only GWAS significant SNPs was not conducted, as a 2 SNP score (after linkage disequilibrium (LD)-based pruning) would be underpowered. LD pruning was performed via PLINK on a random sample of 10,000 individuals using an r^2^>0.05 in a 250kb window. The SNP with the lowest p-value was selected from each of the LD-clumped SNP sets. Where 2 or more SNPs from a set had the same p-value, the SNP with the larger beta coefficient was used. The scores were calculated in PLINK to produce a per-allele weighted score (without mean imputation). Using STATA, deciles of scores were computed and modelling the effect of the PRS on risk was adjusted for age, sex, chip and PCs 1-8.

### Data mining

SNPs associated (at genome-wide significance) with risk-taking behaviour were further investigated for influence on nearby genes (Variant Effect Predictor, VEP ^20^) and for reported associations with relevant traits (GWAS catalogue ^21^). Descriptions and known or predicted functions of implicated genes were compiled (GeneCards www.genecards.org and Entrez Gene www.ncbi.nlm.nih.gov/entrez/query.fcgi?db=gene) and global patterns of tissue expression were assessed (GTEx ^22^). Exploratory analyses of the impact of significant loci on the expression of nearby genes were carried out using the GTEx Portal “Test your own eQTL” function ^22^. In the 13 brain regions available in the GTEx dataset, we tested for associations between rs13084531 and *CADM2* expression, and between rs9379971 and the expression of *POM121L2*, *PRSS16*, *ZNF204P* and *VN1R10P*.

### SNP heritability and genetic correlation analyses

Linkage Disequilibrium Score Regression (LDSR) ^23^ was applied to the GWAS summary statistics to estimate the risk-taking SNP heritability (h^2^_SNP_). LDSR was also used to assess genetic correlations between risk-taking behaviour and relevant psychiatric, cognitive and behavioural traits, namely: ADHD, schizophrenia, BD, MDD, anxiety, PTSD, smoking status (ever smoked), lifetime cannabis use, fluid intelligence, years of education, obesity and alcohol use disorder.

The importance of the brain in regulation of obesity has been demonstrated ^24^, with reward circuits being implicated. The prevalence of obesity in psychiatric illness and the possibility of over-eating being a problem behaviour suggest that there might be a connection between obesity and risk-240 taking behaviour. Thus two measures of obesity were included: Body-Mass Index (BMI) as a measure of total obesity ^24^ and waist-to-hip ratio adjusted for BMI (WHRadjBMI), reflecting metabolically-detrimental central obesity ^25^.

For the ADHD, schizophrenia, BD, MDD, anxiety, PTSD, and smoking status, we used GWAS summary statistics provided by the Psychiatric Genomics Consortium (http://www.med.unc.edu/pgc/) ^26–32^. For the two obesity phenotypes, GWAS summary statistics for BMI ^24^ and WHRadjBMI ^25^ were taken from the consortium for the Genetic Investigation of Anthropometric Traits (http://portals.broadinstitute.org/collaboration/giant). Summary statistics for years of education ^33^ and fluid intelligence ^34^ were downloaded as instructed in the respective publications. Summary statistics for the GWAS of lifetime cannabis use were provided by the International Cannabis Consortium ^35^. Summary statistics for GWAS of alcohol consumption ^36^ and brain structure volumes ^37^ were provided by the authors. Alcohol use disorder was defined using DSM-5 criteria ^38^. For this phenotype, a GWAS meta-analysis on genotypes imputed to 1000 Genomes was run with five datasets: COGEND, COGEND2, COGEND-23andMe, COGA, and FSCD. In total there were N=2 983 cases with alcohol use disorder and N=1 169 controls. Descriptions of the datasets are in the Supplementary information.

## Results

### Demographic characteristics

A subset of 20 335 participants had repeated assessment of risk-taking behaviour. Reproducibility was good, with consistent responses in 81% of all participants (inconsistent 13%, missing 6%, Supplementary Table 2). Participants with probable mood disorders ^39,40^ showed comparable reproducibility compared to those without (consistent 80% vs 82%, inconsistent 15% vs 12%, missing 5% vs 5%, respectively).

For all analyses (discovery, replication and PGRS), small but consistent differences were observed between controls and risk-takers with regard to age and BMI (Table 1), but striking differences were observed for sex distribution, smoking and history of mood disorders: risk-takers (compared to non risk-takers) were more often men, more likely to be current or ever-smokers and more likely to suffer from depression, report an addiction or to have used cannabis. Risk-takers were also more likely to have a university/college degree.

### GWAS of risk-taking behaviour

GWAS results for risk-taking are summarised in Figure 1 (Manhattan plot), Figure 1 inset (QQ plot) and Supplementary Table 3. The GWAS data test statistics showed modest deviation from the null (λ_GC_ =1.13). Considering the sample size, the deviation was negligible (λ_GC_ 1000=1.002). LDSR suggested that deviation from the null was due to a polygenic architecture in which h^2^_SNP_ accounted for approximately 4% of the population variance in risk-taking behaviour (observed scale *h*^2^_SNP_=0.058 (SE 0.006)), rather than inflation due to unconstrained population structure (LD regression intercept=1.003 (SE 0.008)).

**Figure 1:**
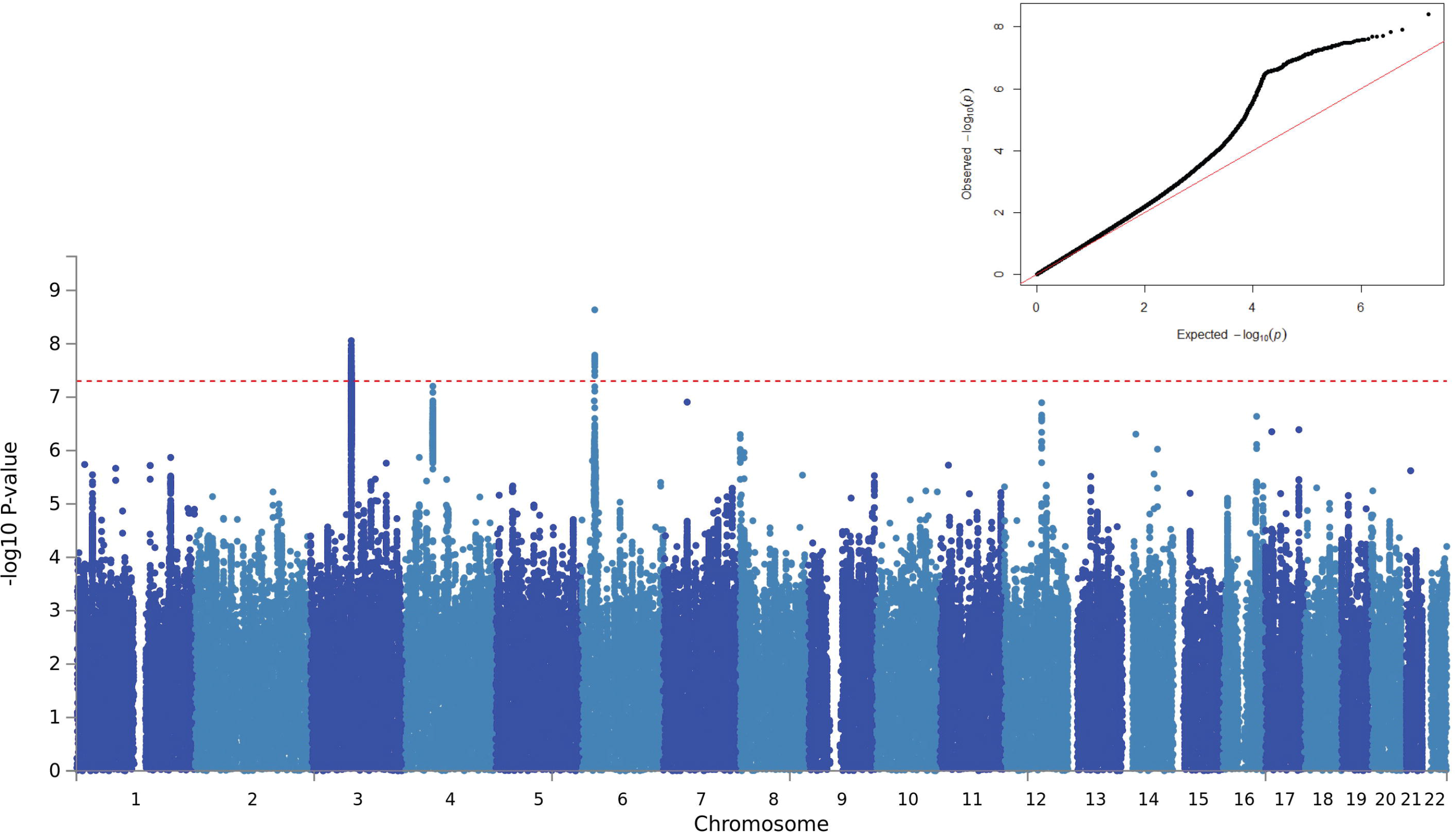
Results of a genome-wide association study of self-reported risk-taking behaviour (1000 Genome imputation). SNPs are plotted along the X axis by chromosome and position, with strength of association with self-reported risk-taking behaviour plotted on the Y axis. The red line indicates the threshold for GWAS significance (P≤5e^−8^). Inset: QQ plot demonstrates deviation from null expectation (solid red line) of the GWAS results (black data points).

Two loci were associated with risk-taking behaviour at genome-wide significance, on chromosome 3 and chromosome 6 (Figure 1 and Supplementary Table 3). The index SNP on chromosome (chr) 3, rs13084531, lies within the *CADM2* gene, however linkage disequilibrium (LD) suggests that the signal also encompasses *miR5688*, and borders a *CADM2* anti-sense transcript (*CADM2-AS2*, Figure 2a). The minor allele of rs13084531 was associated with increased risk-taking (G allele, MAF 0.23, Odds Ratio (OR) 1.07, Confidence interval (CI) 1.04-1.09, P 8.75×10^−9^). Conditional analysis of the chr3 locus (including rs13084531 as a covariate) is suggestive of a second signal (index SNP rs62250716, MAF 0.36, OR 0.96, CI 0.94−0.98, P 8.53=10^−5^, LD r^2^=0.16 with rs13084531, Figure 2b and Supplementary Table 3). The LD structure across the chr3 locus supports the possibility of two distinct signals (Supplementary Figure 1).

**Figure 2:**
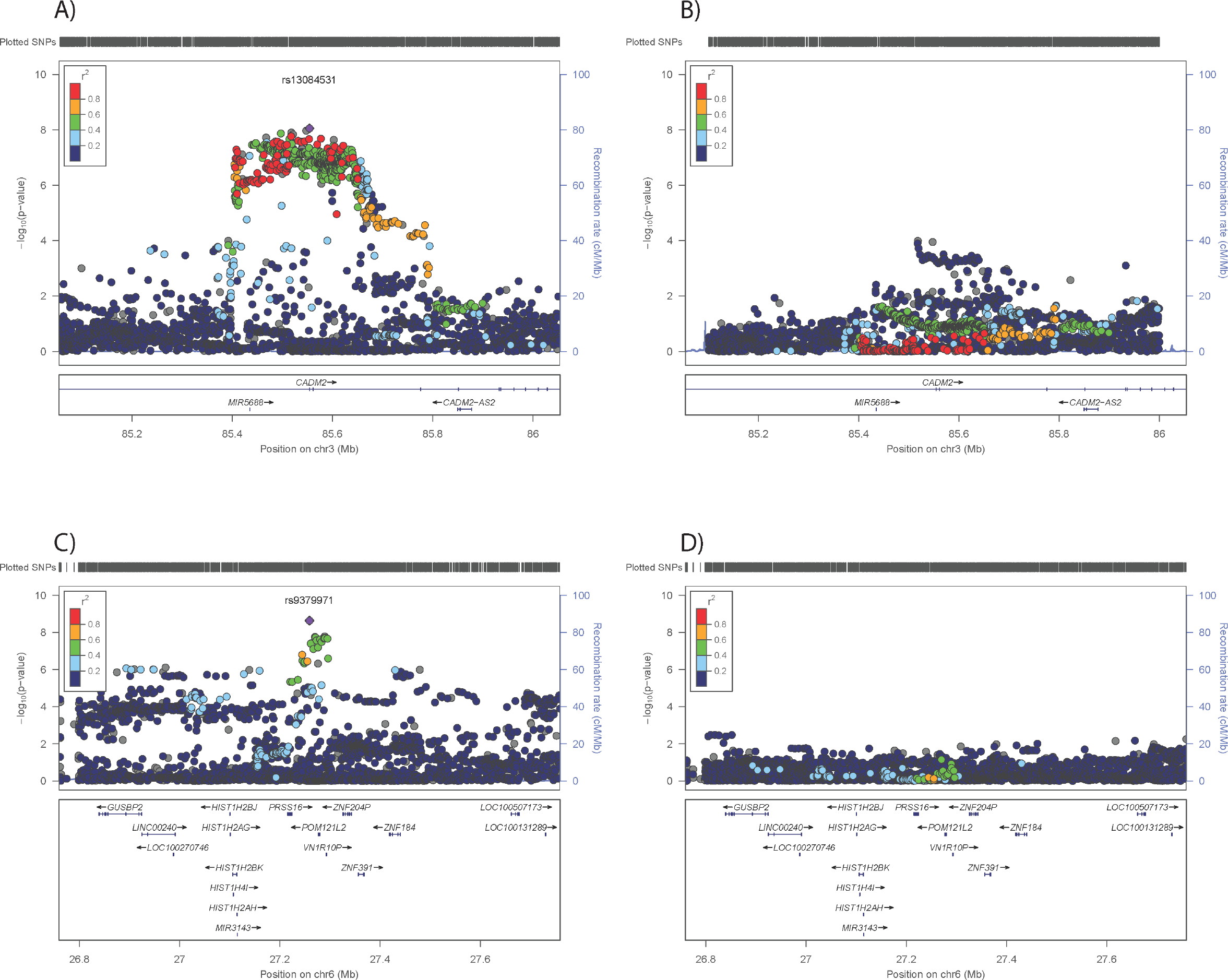
Regional plots for risk-taking-associated loci. A) Chr3 main analysis results; B) results of analysis conditioned on Chr3 rs13084531; C) Chr6 main analysis results; D) results of analysis conditioned on Chr6 rs9379971. The index SNP is shown as a purple diamond.

The chr6 locus lies within the gene-rich HLA region (Figure 2c), where index SNP rs9379971 demonstrated an association between the minor allele and decreased risk-taking (A allele, MAF 0.35, OR 0.95, CI 0.93-0.97, P 2.31×10^−9^). Conditional analysis (including rs9379971 as a covariate) and assessment of the LD structure across this locus indicated that the associated region probably includes only one signal (Figure 2d, Supplementary Table 3 and Supplementary Figure 2).

Rerunning the GWAS with the second genetic data release (Supplementary Figure 3) gave similar results, with a modest deviation from the null (λ_GC_ =1.10, adjusted for sample size λ_GC_ 1000= 1.002). Consistent with the 1000Genomes analysis, LDSR suggested that deviation from the null was due to a polygenic architecture with h^2^_SNP_ accounting for approximately 5% of the population variance in risk-taking behaviour (observed scale *h*^2^_SNP_=0.055 (SE 0.006). The same *CADM2* locus was GWAS significant (rs62250713, beta 0.0614, se 0.01, p= 8.289×10^−10^, minor allele A, MAF 0.36) but the locus on chromosome 6 did not meet the threshold for significance.

### Replication analysis

Both the *CADM2* and chr6 loci demonstrated significant (p<0.025) associations with risk-taking behaviour in the replication analyses (Supplementary Table 4). The *CADM2* locus demonstrated effect sizes comparable with those for the discovery analysis (rs13084531 beta 0.067 for discovery and beta 0.054 se 0.011 replication). In contrast, the Chr6 locus demonstrated 2 to 6-fold weaker effects (rs9379971, discovery beta −0.063, replication beta −0.010). The *CADM2* locus met the threshold for GWAS significance in the meta-analysis (Supplementary Table 4) but the Chr6 locus did not. The significant p value for heterogeneity suggests that this association is a false-positive finding.

### PRS analysis

The PRS were significant predictors of risk-taking behaviour, at all p thresholds and the variance explained by the model including the PRS was between 0.034 (PRS p<1×10^−5^) and 0.037 (PRS p<0.05) (Supplementary Table 5).

### Data mining

As with the majority of SNPs identified by GWAS, the genome-wide significant SNPs in both loci are non-coding. Current prediction models ascribe only non-coding modifier functions to the 81 genome-wide significant SNPs (VEP ^20^, Supplementary Table 6). Expression quantitative trait analysis directly tests association of the index SNPs with expression of nearby transcripts. The chr3 index SNP (rs13084531) lies within the *CADM2* gene and adjacent to miR5688 and *CADM2-AS2* (Figure 2 and Supplementary Table 7). Currently most miRs are predicted (but not reliably proven) to influence transcription of hundreds or thousands of genes. Furthermore, analysing transcription levels of miRs is challenging. Similarly, the importance of antisense transcripts such as *CADM2-AS2* is unclear and difficult to assess. *CADM2*, which encodes cell adhesion molecule 2 (also known as synaptic cell adhesion molecule, SynCAM2), is a plausible target gene as it is predominantly expressed in the brain (Supplementary Figure 4 A). The risk allele at rs13084531 was associated with increased *CADM2* mRNA levels in several regions of the brain (including the caudate basal ganglia and putamen basal ganglia, hippocampus and hypothalamus, Supplemental Figure 5). *CADM1*, a related cell adhesion molecule, demonstrates overlapping and co-regulated (albeit inversely) expression patterns ^41^. It is worth noting that *CADM1* shows a similar, albeit less brain-specific, expression pattern (Supplementary Figure 4 B) and that genetic deletion of *Cadm1* in mice results in behavioural abnormalities, including anxiety ^42^.

Excitement-seeking is a behavioural trait closely related to risk-taking behaviour ^43^, however the locus reported for excitement-seeking was non-significant in this study (Chr2, rs11126769, LD R^2^ with the reported rs7600563= 0.862, major T allele, Beta 0.016, se 0.011, p=0.1167). Other potentially problematic behaviours which can be related to risk-taking propensity have identified the *CADM2* locus (Supplemental Table 8): A recent GWAS of alcohol consumption ^44^ identified a significant signal in the *CADM2* locus, where the G allele of rs9841829 was associated with increased alcohol consumption. The same SNP demonstrates genome-wide significance with increased risk-taking behaviour in this study (G, Beta 0.0635, se 0.012 p=3.34×10^−8^, Supplemental Table 3), whilst conditional analysis (Supplemental Table 3) indicates that the signal for alcohol consumption and risk-taking is the same. A GWAS of lifetime cannabis use also highlighted the *CADM2* locus (gene-based rather than SNP-based) ^35^. Cognitive function plays a role in traits such as risk-taking, therefore it is worth noting that a GWAS of executive functioning and information processing speed in non-demented older adults from the CHARGE (Cohorts for Heart and Aging Research in Genomic Epidemiology) consortium found that genetic variation in the *CADM2* gene was associated with individual differences in information processing speed ^45^. The allele of rs17518584 (LD r^2^=0.45 with rs13084531, LD r^2^=0.34 with rs62250716) associated with increased processing speed was associated with reduced (self-reported) risk-taking in the current study (Supplementary Table 8, p=1.17×10^−7^). Furthermore, a GWAS of educational attainment in the UK Biobank cohort demonstrated a significant signal in *CADM2* ^46^. The effect allele of rs56262138 (LD r^2^=0.00 with rs13084531, LD r^2^=0.00 with rs62250716) for increased educational attainment showed a negative effect on risk-taking behaviour (Supplementary Table 8, p=0.0210).

Day *et al* reported an association between the *CADM2* locus and age of reproductive onset in UK Biobank. In a secondary analysis, they also report an association between the same locus, *CADM2*, and risk-taking behaviour (the same phenotype as was used here). However, differences in quality control procedures mean that the lead SNP reported by Day *et al* was not available in our analysis. During the revision of this paper, Boutwell *et al* ^10^ replicated the association between the *CADM2* locus and a number of personality traits including risk-taking (“do you feel comfortable or uncomfortable with taking risks?”), in an independent data set (n∽140,000).

The *CADM2* locus has also been tentatively associated with longevity ^47^ (Supplementary Table 8) rs9841144, LD r^2^=0.99 with rs13084531, LD r^2^=0.16 with rs62250716), but associations between CADM2 SNPs and longevity, survival and attaining 100 years of age in that study were inconsistent, limiting the interpretation of these signals in the context of risk-taking behaviour.

### Genetic correlations

Looking up the risk-taking SNPs in the GWAS results of psychiatric conditions demonstrated little or no effect of the CADM2 SNPs in ADHD, SCZ, PTSD, BPD or MDD (Supplementary Table 9). In contrast, when considering the entire genome we found significant positive genetic correlations between the risk-taking phenotype and ADHD (r_g_=0.31, SE=0.13, p=0.01), schizophrenia (r_g_=0.27, SE=0.04, p=4.54x10^−11^), BD (r_g_=0.26, SE=0.07, p=1.73×10^−4^), PTSD (r_g_=0.51, SE=0.17, p=0.0018), lifetime cannabis use (r_g_=0.41, SE=0.11, p=0.0001) and smoking (r_g_=0.17, SE=0.07, p=0.01) and a negative genetic correlation with fluid intelligence (r_g_=−0.15, SE=0.05, p=0.0013, Table 2). We found no significant genetic correlation between risk-taking and MDD, anxiety or years of education (Table 2). There was also a significant genetic correlation between risk-taking and BMI (r_g_=0.10, SE=0.03, p=0.003), but a similar correlation was not found for WHRadjBMI. The non-significant genetic correlation with alcohol use disorder was interesting because of the strength of the coefficient (r_g_=0.22, SE0.31, p=0.47), however was likely underpowered due to the modest size of the GWAS (n=4 171) and we draw no conclusions about this correlation.

**Table 2.** Genetic correlation between risk-taking and traits relevant to psychiatric disorders.

| Phenotype | rg | se | p |
| --- | --- | --- | --- |
| ADHD | 0.378 | 0.054 | **1.80x10<sup>-12</sup>** |
| SCZ | 0.265 | 0.040 | **4.54x10<sup>-11</sup>** |
| BD | 0.261 | 0.070 | **1.73x10<sup>-4</sup>** |
| MDD | 0.069 | 0.084 | 0.4120 |
| PTSD | 0.513 | 0.165 | **0.0018** |
| Anxiety (case control) | -0.090 | 0.132 | 0.4963 |
| Anxiety (quantitative) | -0.123 | 0.156 | 0.4289 |
| Ever smoker | 0.174 | 0.068 | **0.0102** |
| Alcohol (heavy vs light) | 0.249 | 0.234 | 0.2873 |
| Alcohol (quantitative) | 0.248 | 0.082 | **0.0026** |
| Alcohol use disorder | 0.221 | 0.306 | 0.4700 |
| Lifetime cannabis use | 0.406 | 0.107 | **1.00x10<sup>-4</sup>** |
| Caudate volume | 0.049 | 0.078 | 0.5268 |
| Accumbens volumes | 0.195 | 0.143 | 0.1710 |
| Fluid intelligence | -0.151 | 0.047 | **0.0013** |
| Years of education | -0.023 | 0.033 | 0.4873 |
| BMI | 0.102 | 0.034 | **0.0028** |
| WHRadjBMI | 0.087 | 0.047 | 0.0655 |

Where: MDD, major depressive disorder; BD, bipolar disorder; SCZ, schizophrenia; ADHD, attention deficit hyperactivity disorder; BMI, body mass index; WHRadjBMI; waist:hip ratio adjusted for BMI; Alcohol dependence DSM 5 Criteria; rg, regression coefficient; se, standard error of the regression coefficient; p, pvalue for theregression analysis.

## Discussion

There is a growing emphasis on the importance of using phenotypic traits which cut across traditional diagnostic groups to investigate the biological basis of psychiatric disorders. Risk-taking behaviour is one such trans-nosological characteristic, recognised clinically as a feature of several disorders, including ADHD, SCZ and BD. In this study we identified two loci, on 3p12.1 and 6p22.1, that were associated with self-reported risk-taking behaviour. Replication in an independent set of samples and meta-analysis confirmed the association between risk-taking behaviour and the *CADM2* locus on Chr3 but not the Chr6 locus. The PRS were significant predictors of risk-taking behaviour in a further independent sample set.

The chr6 locus falls within the HLA region which encodes a large number of genes and is extremely complicated genetically. The false positive association detected could be because the first data release were selected based on (extremes of) lung function measurements ^48^. Considering the potential inflammatory component of lung function and the role of the HLA region in inflammatory responses, it is perhaps not surprising that the discovery analysis demonstrated stronger effect sizes for this locus than the randomly selected general population samples included in the replication analysis.

A key finding of our study was the positive association between Chr3 SNP, rs13084531, and risk-taking behaviour as well as *CADM2* expression levels. Here, the allele associated with increased self-reported risk-taking behaviour was also associated with increased *CADM2* expression. It is of interest that lack of *Cadm1* in mice was associated with anxiety-related behaviour ^42^ and that both *CADM1* and *CADM2* were identified as BMI-associated loci ^24^ suggesting that *CADM2* and related family members may be involved in balancing appetitive and avoidant behaviours.

Day and colleagues recently identified 38 genome-wide significant loci for age at first sexual intercourse within the UK Biobank cohort ^2^ and two of these loci were within the 3p12.1 region, close to *CADM2* (rs12714592 and rs57401290). The association between rs57401290 (and SNPs in LD) and age at first sexual intercourse was also observed for a number of behavioural traits, including number of sexual partners, number of children and risk-taking propensity (the same phenotype as was used in this study). In addition, *CADM2* also showed association with information processing speed ^45^ and educational attainment ^46^, highlighting the complexity of relationships between cognitive performance and risk-taking. Taken together, this evidence suggests that *CADM2* plays a fundamental role in risk-taking behaviours, and may be a gene involved in the nexus of cognitive and reward-related processes that underlie them.

A perhaps surprising observation was the increased frequency of having a university degree in self-reported risk-takers, compared to controls, despite the negative (albeit non-significant) association between years of education and risk-taking behaviour. It is important to note that risk-taking behaviour includes a number of different aspects, including delayed gratification, assessment of positive and negative consequences of risk, impulse control, reward signalling. It is possible that risk-taking behaviour assessed in a clinical mental health setting could reflect a different aspect of these processes compared to self-reported risk-taking behaviour. Risk-taking behaviour assessed in a clinical mental health setting might demonstrate significantly different associations with education, compared to self-reported risk-taking behaviour. These observations underscore the complexity between risk-taking and educational attainment, and highlight differences between genetic and phenotypic relationships. They may also be indicative of selection bias within the UK Biobank cohort towards more highly educated individuals.

Another key finding was genetic correlation between self-reported risk-taking and obesity. Although there are likely to be a range of potential mechanisms linking risk-taking behaviour with obesity, evidence of a shared genetic component is in keeping with work that has highlighted the importance of the central nervous system in the regulation of obesity (BMI), particularly brain regions involved in cognition, learning and reward ^24^. In contrast, central fat accumulation (WHRadjBMI) is primarily regulated by adipose tissue ^25^ which fits with the lower, non-significant genetic correlation between risk-taking behaviour and this measure. Two SNPs (rs13078807 and rs13078960) in the *CADM2* locus have previously been associated with BMI ^24, 49, 50^, but whilst these SNPs tag each other (LD r^2^=0.99), the LD between the risk-taking index SNP or possible secondary signal is low (LD r^2^=0.31 and 0.01 for rs13084531 and rs62250716 respectively), suggesting that these are distinct signals.

It is perhaps unsurprising that we identified genetic correlations between risk-taking and smoking. Similarly, risk-taking and impulsive behaviour is a core feature of ADHD and BD, suggesting substantial genetic overlap between variants predisposing to risk-taking behaviour and these disorders. The genetic correlation between risk-taking and schizophrenia is of interest because schizophrenia is commonly comorbid with substance abuse disorders ^51^. The correlation between risk-taking and PTSD is perhaps plausible if we accept that risk-takers may be more likely to find themselves in high-risk situations with the potential to cause psychological trauma. Overall, these correlations suggest that studying dimensional traits such as risk-taking has the potential to inform the biology of complex psychiatric disorders.

### Strengths and limitations

We acknowledge that Day *et al* have previously reported an association for risk-taking within the *CADM2* locus. Strengths of our study include the use of a more conservative and standardised methodology and reporting of results across the entire genome. A risk-taking locus was identified in the *CADM2* locus and we have shown that *CADM2* may contain a second signal. Furthermore, we have investigated the possibility of a sex-specific effect of these loci, provided evidence highlighting possible candidate genes at both loci and confirmed the importance of this phenotype in relation to psychiatric illness. In short, our report provides a fuller understanding of the genetic basis of risk-taking behaviour. Despite this, we highlight some limitations. The risk-taking phenotype used was a self-reported measure, based on response to a single question, and is therefore open to responder bias. It is also plausible that there are distinct subtypes of risk-taking behaviour (for example disinhibition, sensation-seeking and calculated risks). Whether the single question used in our analyses captures all, or only some, of these is not clear. Having identified genetic loci associated with other traits related to risk-taking and other problem behaviours (such as alcohol consumption and cannabis use) provides added support for the validity of this phenotype. It would be of interest to investigate whether the loci identified here are also associated with more quantitative and objective measures of risk taking; however, such measures were not available in the UK Biobank dataset.

### Conclusion

In summary, we have identified a polygenic basis for self-reported risk-taking behaviour and the *CADM2* locus which contains variants likely to play a role in predisposition to this complex but important phenotype. The identification of significant genetic correlations between risk-taking and several psychiatric disorders, as well as with smoking and obesity, suggest that future work on this trait may clarify mechanisms underlying several common psychopathological and physical health conditions, which are important for public health and wellbeing.

## Acknowledgements

This research was conducted using the UK Biobank resource. UK Biobank was established by the Wellcome Trust, Medical Research Council, Department of Health, Scottish Government and Northwest Regional Development Agency. UK Biobank has also had funding from the Welsh Assembly Government and the British Heart Foundation. Data collection was funded by UK Biobank. JW is supported by the JMAS Sim Fellowship for depression research from the Royal College of Physicians of Edinburgh (173558). AF is supported by an MRC Doctoral Training Programme Studentship at the University of Glasgow (MR/K501335/1). DJS acknowledges the support of the Brain and Behaviour Research Foundation (Independent Investigator Award 1930) and a Lister Prize Fellowship (173096). EMT is supported by a University Research Fellowship (UF140705) from the Royal Society. JC acknowledges the support of The Sackler Trust and is part of the Wellcome Trust funded Neuroimmunology of Mood and Alzheimer’s consortium that includes collaboration with GSK, Lundbeck, Pfizer and Janssen & Janssen. The work at Cardiff University was funded by Medical Research Council (MRC) Centre (G0800509) and Program Grants (G0801418). The funders had no role in the design or analysis of this study, decision to publish, or preparation of the manuscript.

## Conflict of interest

JPP is a member of UK Biobank advisory committee; this had no bearing on the study. No other conflicts of interest.

**Supplementary information is available at Molecular Psychiatry’s website.**

